# A Single Adaptive Mutation in Sodium Taurocholate Cotransporting Polypeptide Induced by Hepadnaviruses Determines Virus Species-specificity

**DOI:** 10.1101/391490

**Authors:** Junko S Takeuchi, Kento Fukano, Masashi Iwamoto, Senko Tsukuda, Ryosuke Suzuki, Hideki Aizaki, Masamichi Muramatsu, Takaji Wakita, Camille Sureau, Koichi Watashi

## Abstract

Hepatitis B virus (HBV) and its hepadnavirus relatives infect a wide range of vertebrates from fish to human. Hepadnaviruses and their hosts have a long history of acquiring adaptive mutations. However, there are no reports providing direct molecular evidence for such a coevolutionary “arms race” between hepadnaviruses and their hosts. Here, we present evidence suggesting the adaptive evolution of the sodium taurocholate cotransporting polypeptide (NTCP), an HBV receptor, has been influenced by virus infection. Evolutionary analysis of the NTCP-encoding genes from 20 mammals showed that most NTCP residues are highly conserved among species, exhibiting evolution under negative selection (dN/dS < 1); this observation implies that the evolution of NTCP is restricted by maintaining its original protein function. However, 0.7 % of NTCP amino acid (aa) residues exhibit rapid evolution under positive selection (dN/dS > 1). Notably, a substitution at aa 158, a positively selected residue, converting the human NTCP to a monkey-type sequence abrogated the capacity to support HBV infection; conversely, a substitution at this residue converting the monkey Ntcp to the human sequence was sufficient to confer HBV susceptibility. Together, these observations suggested that positive selection at aa 158 was induced by virus infection. Moreover, the aa 158 sequence determined attachment of the HBV envelope protein to host cell, demonstrating the mechanism whereby HBV infection would create positive selection at this residue in NTCP. In summary, we provide the first evidence in agreement with the function of hepadnavirus as a driver for inducing an adaptive mutation in host receptor.

**Importance:** Hepatitis B virus (HBV) and its hepadnavirus relatives infect a wide range of vertebrates, with a long infectious history (hundreds of millions of years). Such a long history generally allows adaptive mutations in hosts to escape from infection, while simultaneously allowing adaptive mutations in viruses to overcome host barriers. However, there is no published molecular evidence for such a coevolutionary “arms race” between hepadnaviruses and hosts. In the present study, we performed coevolutionary phylogenetic analysis between hepadnaviruses and the sodium taurocholate cotransporting polypeptide (NTCP), an HBV receptor, combined with virological experimental assays for investigating the biological significance of NTCP sequence variation. Our data provide the first molecular evidences supporting that HBV-related hepadnaviruses drive adaptive evolution in the NTCP sequence, including a mechanistic explanation of how NTCP mutations determine host viral susceptibility. Our novel insights enhance our understanding of how hepadnaviruses evolved with their hosts, permitting the acquisition of strong species-specificity.

## Introduction

Over 250 million people are chronically infected worldwide with hepatitis B virus (HBV), and hepatitis B resulted in 887,000 deaths in 2015 (1). HBV and its hepadnavirus relatives infect a wide range of vertebrates, including fish, amphibians, reptiles, birds, and mammals (2-10). Phylogenetic analyses suggest that the coevolutionary history of hepadnaviruses and their hosts spans hundreds of millions of years (2, 6, 7, 9). During the long-term coevolution of virus and host, host proteins that interact directly with a virus accumulate adaptive mutations to permit escape from virus infection, while viruses acquire mutations that overcome host barriers (11). In the course of this coevolutionary “arms race”, viruses continuously exert selective pressures on their host, thereby inducing multiple adaptive mutations in the host genome (12). One of the best-known examples of such coevolution is embodied by the conflict between (host-encoded) restriction factors and their (virus-encoded) viral antagonists (11). Host restriction factors inhibit viral replication at different steps during virus proliferation in the host. Conversely, viruses encode factors that antagonize the function of host restriction factors and thereby circumvent the host’s attempt to eliminate viruses (13). During the course of long-term viral selective pressures, some host restriction factors evolve rapidly, a process that is evidenced as a ratio of nonsynonymous to synonymous mutations in the host gene (the dN/dS ratio) that exceeds 1 (“positive selection”) (14). For example, host restriction factors against human immunodeficiency virus type 1 (HIV-1), including tripartite motif-containing protein 5-alpha (TRIM5α) (15), apolipoprotein B mRNA editing enzyme catalytic polypeptide-like 3G (APOBEC3G) (16), bone marrow stromal antigen 2 (BST2, also known as tetherin, CD317, and HM1.24) (17-20), and SAM domain and HD domain 1 (SAMHD1) (21, 22), have been reported to exhibit rapid evolution (dN/dS > 1), likely due to the selective pressure exerted by HIV-1 infection. Regarding the coevolution of hepadnaviruses and host restriction factors, Abdul *et al.* recently reported an evolutionary analysis of an HBV restriction factor, the Structural Maintenance of Chromosomes (Smc) 5/6 complex (23), a complex originally identified based on its housekeeping function in genomic stability (24). However, Abdul *et al.* did not detect a clear signature of positive selection that was suggested to be induced by hepadnavirus infection. In contrast, Enard *et al.* reported that host proteins interacting with viruses with a long history display higher rates of adaptive mutations (12); those authors showed that host proteins reported to interact with HBV exhibited a strong signature of adaptation during a coevolution with viruses, which was in a similar degree to that seen for HIV-1-interacting host proteins. However, molecules subject to such a selective pressure by hepadnaviruses have not (to our knowledge) been identified to date.

Hepadnaviruses infect their hosts in a highly species-specific manner; for instance HBV can infect only humans, chimpanzees, and treeshrews, but not monkeys, including both Old-World and New-World monkeys (25). The sodium taurocholate cotransporting polypeptide [NTCP, also designated as solute carrier family 10A1 (SLC10A1)] was recently identified as a host factor that functions as an HBV entry receptor; NTCP, which originally was characterized as a hepatic transporter for the uptake of bile acids by hepatocytes, binds to the HBV envelope protein, notably to the preS1 region, thereby mediating viral entry into the host (26). NTCP has been suggested to be a key determinant of the species-specificity of HBV, as primary monkey hepatocytes can support the replication of intracellular HBV but not the entry of the virus into host cells (27), and complementation the monkey cells with human NTCP (hNTCP) permits HBV entry and thereby the whole infection cycle both in cell culture and *in vivo* (28, 29). These results indicate that the inability of monkey Ntcp to support HBV infection serves as the species barrier preventing HBV infection in monkey. However, the evolutionary relationship between NTCP sequences in different species and susceptibility to hepadnavirus infection has not been analyzed previously. Virus entry receptors generally have their own original function in cellular physiology. Thus their sequences typically are conserved during evolution to maintain their functional profile, showing a dN/dS < 1 that indicates negative selection (30). Indeed, transferrin receptor (TfR1), which serves as an entry receptor for arenaviruses and mouse mammary tumor virus (31), exhibit such evolutionary negative selection across their entire sequences. Interestingly, however, a small percentage of positions in the TfR1 coding sequence is rapidly evolving under positive selection, and these sites were shown to correspond to a virus-binding surface (31). In another example, the Niemann-Pick disease, type C1 (NPC1), a receptor for filovirus, shows a similar evolutionary property (32). These data strongly suggest that virus infection serve as a pressure for the evolution of viral entry receptors. In addition, positive selection sites on virus entry receptors can provide critical information, permitting the identification of the receptor domains that interface with the virus. However, to our knowledge, there have been no reports of such an evolutionary analysis for a hepadnavirus receptor.

In this study, we focused on the phylogenetic analyses of hepadnaviruses and NTCP based on molecular evolutionary analyses to detect positive selection sites, and investigated the biological significance of NTCP sequence changes using virological experimental assays in our cell culture model (33). Evolutionary analyses of the NTCPs from 20 mammalian species revealed that NTCP was highly conserved among these mammals, but interestingly identified several sites that were subject to positive selection. Among these positive selection sites, the amino acid (aa) at NTCP residue 158 determined the HBV host species-specificity: Our virological experiments showed that a single substitution converting aa 158 of human NTCP to the corresponding aa of cynomolgus macaque Ntcp completely negated the protein’s ability to mediate viral infection, whereas a cynomolgus macaque Ntcp protein mutated to carry the hNTCP aa 158 rendered macaque Ntcp susceptible to the virus, suggesting that this site regulates viral susceptibility. Moreover, aa 158 of NTCP was shown to mediate viral attachment to the host cell surface. In a manner analogous to TfR1 and NPC1, these data suggest that NTCP receives an evolutionary pressure by virus infection, which provides the first evidence in a molecular level for coevolution between hepadnaviruses and hosts.

## Results

### A small portion of sites in *NTCP* evolves under positive selection in mammals

We performed maximum likelihood-based molecular evolutionary analyses (Figure 1) using 20 *NTCP* coding sequences from mammalian species, including human, monkeys, bats, rabbit, rat, and mouse (Table 1). To detect positive selection sites in mammalian *NTCP* s, two pairs of site-models were conducted using PAML ver. 4.8 (34, 35). The differences between neutral models and selection models (M1a vs. M2a, respectively or M7 vs. M8, respectively) were significant (p < 0.01 or p < 0.005, respectively), suggesting that *NTCP* genes have been evolving under positive selection in mammals (Table 2). The M2a analysis estimated that 66.2 % (p0 = 0.662) of the codons were under negative selection with dN/dS ratio (ω0) = 0.086; 33.1 % (p1 = 0.331) were under neutral evolution; and 0.7 % (p2 = 0.007) were under positive selection with dN/dS ratio (ω2) = 4.526. Thus, only a small number of sites in mammalian NTCPs was under positive selection. The same data set also was analyzed to detect positive selection sites by Fixed Effects Likelihood (FEL), Random effects likelihood (REL) (36), and Mixed Effects Model of Evolution (MEME) (37), and 27 positively selected sites were inferred by at least one of the three evolutionary analyses as summarized in Table 3.

**Table 1.** GenBank accession numbers of mammalian NTCP sequences used in this study. ^a^The common name of each mammal is identical to that in Figure 1.

|  | accession # | Scientific name | <sup>a</sup> Common name | Order |
| --- | --- | --- | --- | --- |
| 1 | NM_003049.3 | *Homo sapiens* | Human | Primates (Hominoids) |
| 2 | XM_510035.5 | *Pan troglodytes* | Chimpanzee |  |
| 3 | XM_003824101.2 | *Pan paniscus* | Bonobo |  |
| 4 | XM_002824890.2 | *Pongo abelii* | Sumatran orangutan |  |
| 5 | NM_001283323.1 | *Macaca fascicularis* | Cynomolgus macaque | Primates (Old-World monkeys) |
| 6 | KT382283.1 | *Papio hamadryas* | Hamadryas baboon |  |
| 7 | KT382281.1 | *Chlorocebus aethiops* | Grivet |  |
| 8 | KT326157.1 | *Semnopithecus sp.* | Gray langur |  |
| 9 | KT382285.1 | *Callithrix jacchus* | Marmoset | Primates (New-World monkeys) |
| 10 | KT382284.1 | *Saguinus oedipus* | Cotton-top tamarin |  |
| 11 | KR153327.1 | *Saimiri sciureus* | Squirrel monkey |  |
| 12 | XM_006091030.2 | *Myotis lucifugus* | Little brown bat | Chiroptera |
| 13 | XM_005877950.2 | *Myotis brandtii* | Brandt's bat |  |
| 14 | XM_008138900.1 | *Eptesicus fuscus* | Big brown bat |  |
| 15 | XM_006915181.2 | *Pteropus alecto* | Black flying fox |  |
| 16 | JQ608471.1 | *Tupaia belangeri* | Northern treeshrew | Scandentia |
| 17 | NM_001082768.1 | *Oryctolagus cuniculus* | European rabbit | Lagomorpha |
| 18 | XM_015490579.1 | *Marmota marmota* | Alpine marmot | Rodentia |
| 19 | NM_017047.1 | *Rattus norvegicus* | Brown rat |  |
| 20 | NM_001177561.1 | *Mus musculus* | House mouse |  |

**Table 2.** Log likelihood values and estimated parameters under models in PAML. Two pairs of site-models with CODEML implemented in the PAML ver. 4.8 were performed: the first pair employed M1a (neutral, 2 site classes, ω > 1 not allowed) versus M2a (selection, 3 site classes, ω > 1 allowed), and the second pair consisted of M7 (neutral, 10 site classes, ω > 1 not allowed) versus M8 (selection, 11 site classes, ω > 1 allowed).

| Model | lnL | 2ΔlnL | p value | Estimated parameters | Positive selection sites |
| --- | --- | --- | --- | --- | --- |
| M0 | -5972.703 | | | $\omega = 0.344$ | |
| M1a | -5868.144 | | | $p0 = 0.667, p1 = 0.333$<br>$\omega0 = 0.085, \omega1 = 1$ | |
| M2a | -5863.300 | 9.687 | < 0.01 | $p0 = 0.662, p1 = 0.331, p2 = 0.007$<br>$\omega0 = 0.086, \omega1 = 1, \omega2 = 4.526$ | K157, A351 |
| M7 | -5868.087 | | | $p = 0.237, q = 0.430$ | |
| M8 | -5862.086 | 12.002 | < 0.005 | $p0 = 0.991, p = 0.252, q = 0.469$<br>$p1 = 0.009, \omega = 3.784$ | K157, A351 |

**Figure 1.**
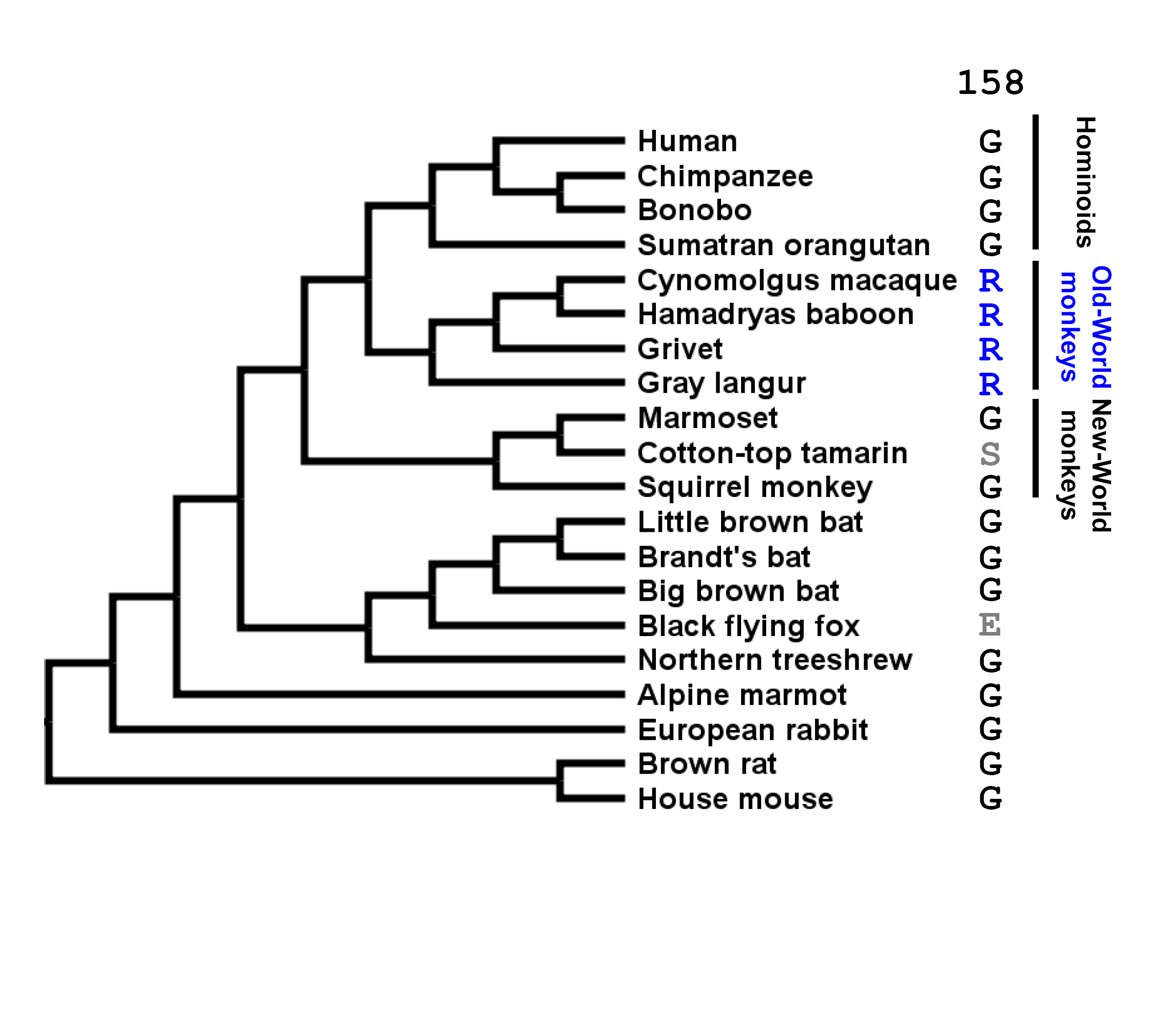
Maximum Likelihood tree of the 20 NTCP sequences analyzed in this study. A Maximum Likelihood (ML) tree of 20 mammalian full-length NTCP sequences was reconstructed using PhyML 3.0 with 1,000 bootstrap resamplings. The corresponding GenBank accession numbers are listed in Table 1. The indicated amino acid sequences correspond to the amino acid at residue 158 in each NTCP species.

### Single positive selection site of NTCP is a key determinant of HBV susceptibility

To date, hepadnaviruses have been found in multiple Hominoids, including chimpanzees and orangutans (38). On the other hand, no HBV or hepadnavirus infections have been reported in Old-World monkeys, even though this family is the closest to the Hominoids (Figure 1) [Although a unique case of HBV genotype D infection was reported in *Macaca fascicularis* (cynomolgus macaques) from Mauritius (39), another recent paper showed that this macaques exhibited no evidence of current or prior HBV infection (29)]. Since Old-World monkeys are the closest species to human that are regarded to be non-susceptible to HBV or hepadnaviruses, we focused on these two lineages for the comparison among *NTCP* genes. Notably, there are 13 aa differences between the human NTCP (NM_003049.3, hNTCP) and the cynomolgus macaque Ntcp (NM_001283323.1, *Macaca fascicularis*, Old-World monkey, mNtcp). Evolutionary analysis comparing the corresponding gene sequences identified five sites (codons 142, 157, 158, 160, and 165) in the *NTCP* open reading frame (ORF) that are under positive selection (Table 3 and Figure 2).

**Table 3.** Positively selected sites inferred under FEL, REL, and MEME. FEL, REL, and MEME were employed to detect the positively selected sites through the DATAMONKEY webserver. ^a^Codons that varied between human (NM_003049.3) and cynomolgus macaque (NM_001283323.1) *NTCP* genes are underlined. ^b,c,d^The following default significant cut-off values were used and are shown in red: p-value < 0.1 for FEL, Bayes factor > 50 for REL, and p-value < 0.1 for MEME.

| <sup>a</sup> site | FEL dN-dS | <sup>b</sup> FEL<br>p-value | REL dN-dS | <sup>c</sup> REL Bayes<br>Factor | MEME $\omega$+ | <sup>d</sup> MEME<br>p-value |
| --- | --- | --- | --- | --- | --- | --- |
| 25 | 3.03 | 0.056 | 0.32 | 9.987 | >100 | 0.034 |
| 62 | 0.575 | 0.278 | -0.094 | 6.955 | >100 | 0.087 |
| 107 | 2.268 | 0.045 | 0.31 | 12.91 | >100 | 0.062 |
| 129 | 2.834 | 0.014 | 1.08 | 37.539 | >100 | 0.022 |
| <u>142</u> | 1.127 | 0.156 | -0.018 | 8.682 | >100 | 0 |
| <u>157</u> | 4.646 | 0.004 | 1.843 | 2566.71 | >100 | 0.007 |
| <u>158</u> | 1.311 | 0.3 | -0.158 | 4.877 | >100 | 0.05 |
| <u>160</u> | 0.959 | 0.437 | -0.21 | 4.018 | >100 | 0.061 |
| <u>165</u> | 1.673 | 0.095 | 0.06 | 9.433 | >100 | 0.071 |
| 166 | 1.152 | 0.067 | 0.188 | 23.465 | >100 | 0.088 |
| 175 | 2.486 | 0.051 | 0.475 | 13.608 | >100 | 0.064 |
| 184 | -1.037 | 0.275 | -0.783 | 0.55 | >100 | 0.038 |
| 198 | 1.592 | 0.099 | 0.031 | 8.999 | >100 | 0.11 |
| 207 | 1.491 | 0.087 | 0.044 | 10.293 | >100 | 0.11 |
| 210 | 0.044 | 0.966 | -0.286 | 3.225 | >100 | 0.047 |
| 223 | 2.671 | 0.051 | 0.22 | 7.959 | >100 | 0.001 |
| 248 | 1.896 | 0.03 | 0.275 | 19.36 | >100 | 0.043 |
| 308 | -0.797 | 0.634 | -0.401 | 1.487 | 25.428 | 0.095 |
| 330 | 1.321 | 0.085 | 0.104 | 13.859 | >100 | 0.11 |
| 333 | 0.883 | 0.094 | 0.175 | 23.698 | >100 | 0.047 |
| 334 | 1.05 | 0.098 | 0.146 | 18.083 | >100 | 0.122 |
| 339 | 1.723 | 0.044 | 0.172 | 15.24 | >100 | 0.062 |
| 342 | 1.03 | 0.169 | -0.022 | 8.713 | >100 | 0.077 |
| 347 | 1.956 | 0.212 | -0.044 | 4.988 | >100 | 0.001 |
| 348 | 0.719 | 0.699 | -0.166 | 2.831 | >100 | 0.008 |
| 351 | 1.961 | 0.705 | 1.346 | 743.714 | >100 | 0.003 |
| 352 | 0.739 | 0.5 | -0.187 | 3.923 | 25.945 | 0.048 |

**Figure 2.**
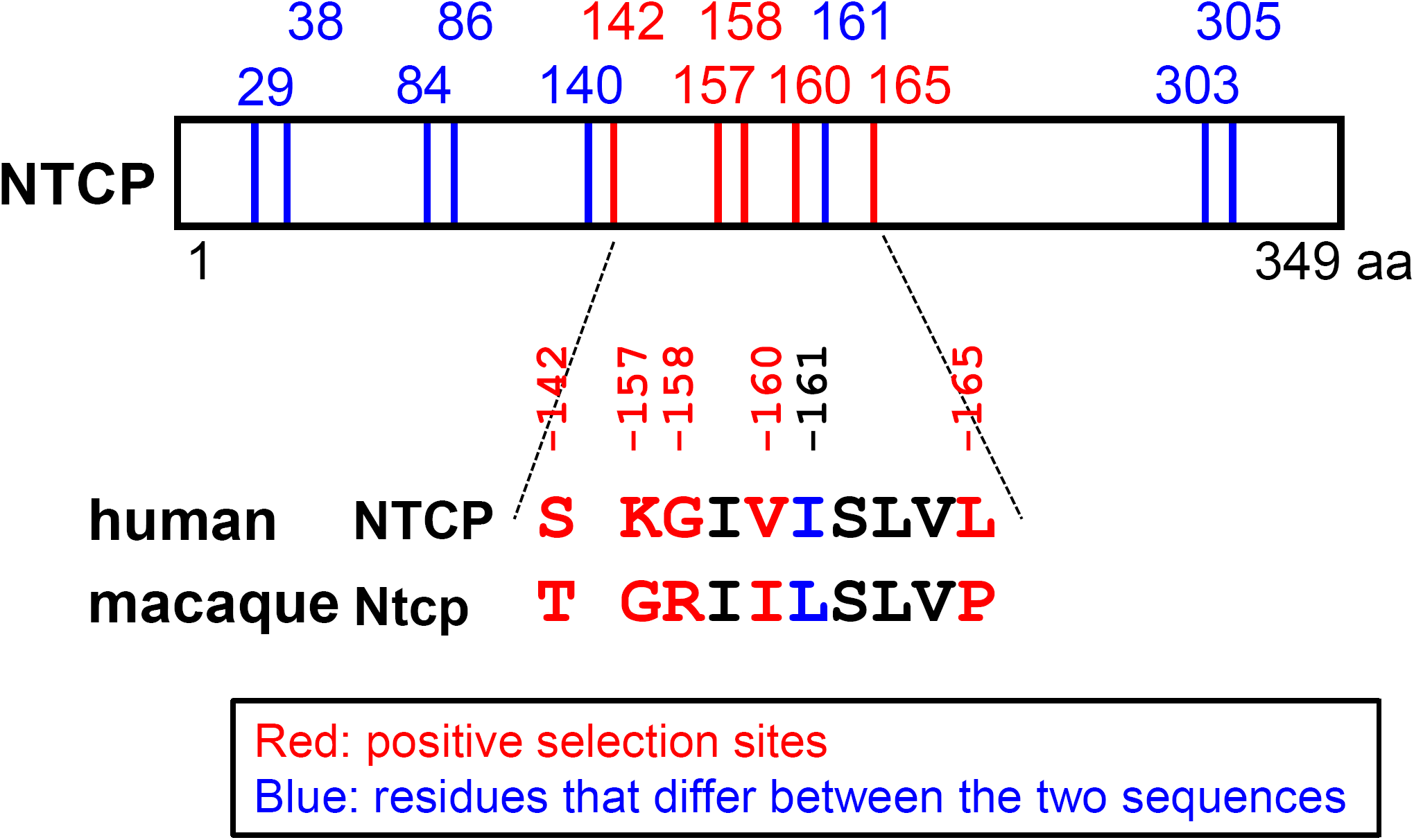
Comparison between human and macaque NTCPs. Schematic representation of the primary amino acid sequence of NTCP. The 13 amino acid positions at which human (NM_003049.3) and cynomolgus macaque (NM_001283323.1) sequences differ are indicated by the numbers above the box showing the NTCP protein and the vertical lines (in red and blue). Among these 13 positions, five residues (indicated in red) were inferred to be under positive selection in the 20 mammalian sequences evaluated in this study. The amino acid sequences at residue 142 and 157-165 are compared between human and cynomolgus macaque, as shown under the box. The indicated numbers correspond to the amino acid positions in human NTCP.

We next addressed whether the amino acids encoded by these sites are involved in susceptibility to HBV infection. We prepared HepG2 cells transiently expressing hNTCP or a series of single-mutant variants in which a single amino acid was replaced by the corresponding macaque residue (S142T, K157G, G158R, V160I, and L165P) (Figure 3A). Similarly, we produced HepG2 cells expressing a variant of hNTCP in which aa 157-165 were replaced by the corresponding macaque sequence [designated hNTCP(m157-165)]; this substitution has been reported to abrogate NTCP’s receptor function (26) (Figure 3A). At 24 h after transfection, protein expression of NTCP or its variants was confirmed by immunoblotting of cell lysates. As shown in Figure 3A, the mutant hNTCPs were expressed at levels equivalent to that of wild-type hNTCP, except for hNTCP-L165P, which had much lower protein expression (Figure 3A). To further validate whether each of the NTCP variants maintained its functional properties as a bile acid transporter, we also measured NTCP-mediated uptake of [^3^H]-taurocholic acid in these cells. All NTCP variants, including hNTCP-L165P, possessed the capacity to uptake bile acid, retaining at least 40 % of the wild-type activity and providing activity 6-to 24-fold that of background (assessed under sodium-free conditions) (Figure 3B). Thus, all of the hNTCP variant-expressing HepG2 cell lines in this experiment retained NTCP functions.

**Figure 3.**
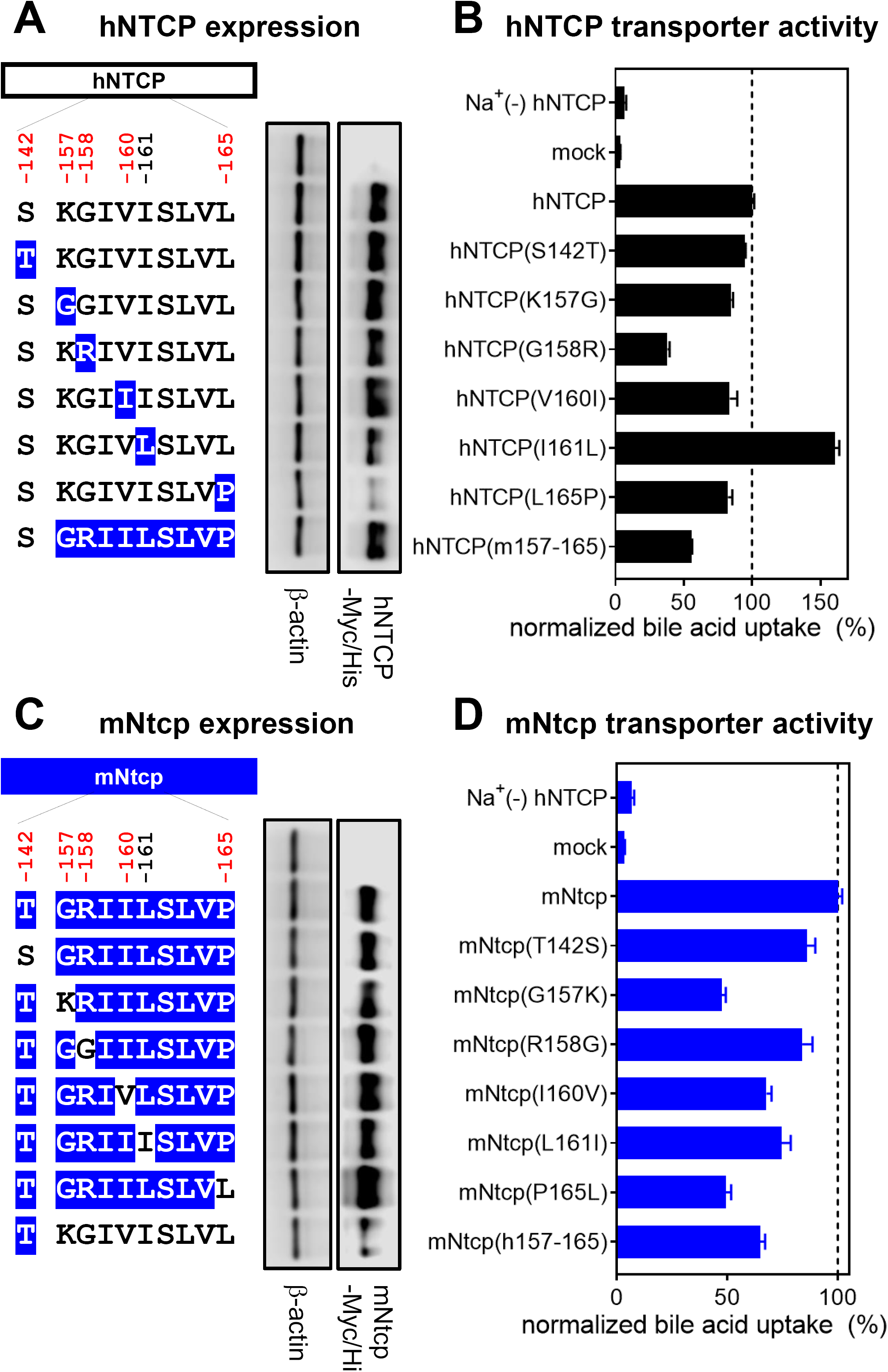
NTCP expression and the transporter activity. (A, C) Left panels, schematic representation of NTCP variants. Black and blue boxes/amino acids indicate those from human and macaque, respectively. The constructs shown in (A) and (C) are variants in which hNTCP (black) and mNtcp (blue) were used as backbones, respectively. Amino acid positions under positive selection are shown in red at the top of the scheme. Right panels, western blotting was used to assess the expression level of Myc/His-tagged NTCP or its variants in HepG2 cells transfected with the respective construct. Actin was used as an internal control. (B, D) Transporter activities of NTCP in these cells were measured by transporter assay using [^3^H]-taurocholic acid as a substrate in either sodium-free buffer (lane 1; to determine background level) or sodium-containing buffer (lanes 2-10). Normalized transporter activities are shown, and were calculated by dividing the detected radioactivity values by NTCP expression levels determined by immunofluorescence analysis. Results are presented as mean ± SEM (n=3).

We then analyzed the ability of these cell lines to support HBV infection. Specifically, an HBV infection assay was performed (per Materials and Methods) (33) to evaluate the cells’ susceptibility to HBV infection by detecting intracellular hepatitis B core antigen (HBcAg) at 13 days post-inoculation by immunofluorescence analysis (Figure 4A, right pictures, red, “DMSO”). The detected levels of HBcAg were normalized to the expression level of NTCP in the respective line, permitting assessment of each NTCP’s ability to support HBV infection (Figure 4A, center graph). In addition, to confirm that the observed fluorescence signals were derived from HBV infection (and did not reflect non-specific background), the HBV infection assay was also performed in parallel in the presence of Myrcludex B (Myr-B), an HBV entry inhibitor (40), a condition that should abrogate the specific signal derived from HBV infection (Figure 4A, right pictures, “Myr-B”). As shown in Figure 4A, HBV infection was observed in the lines expressing wild-type hNTCP, but not in that expressing hNTCP(m157-165) (Figure 4A, lanes 2 and 9), consistent with a previous report (26). HBV infection was supported by all but one of the NTCP variants, hNTCP(S142T), hNTCP(K157G), hNTCP(V160I), hNTCP(I161L), and hNTCP(L165P) (Figure 4A, lanes 3, 4, 6, 7, and 8). In contrast, the level of HBV infection in hNTCP(G158R)-expressing cells was similar to the background level (Figure 4A, lanes 1 and 5), suggesting that the aa 158 was unique among the examined sites in determining HBV susceptibility.

**Figure 4.**
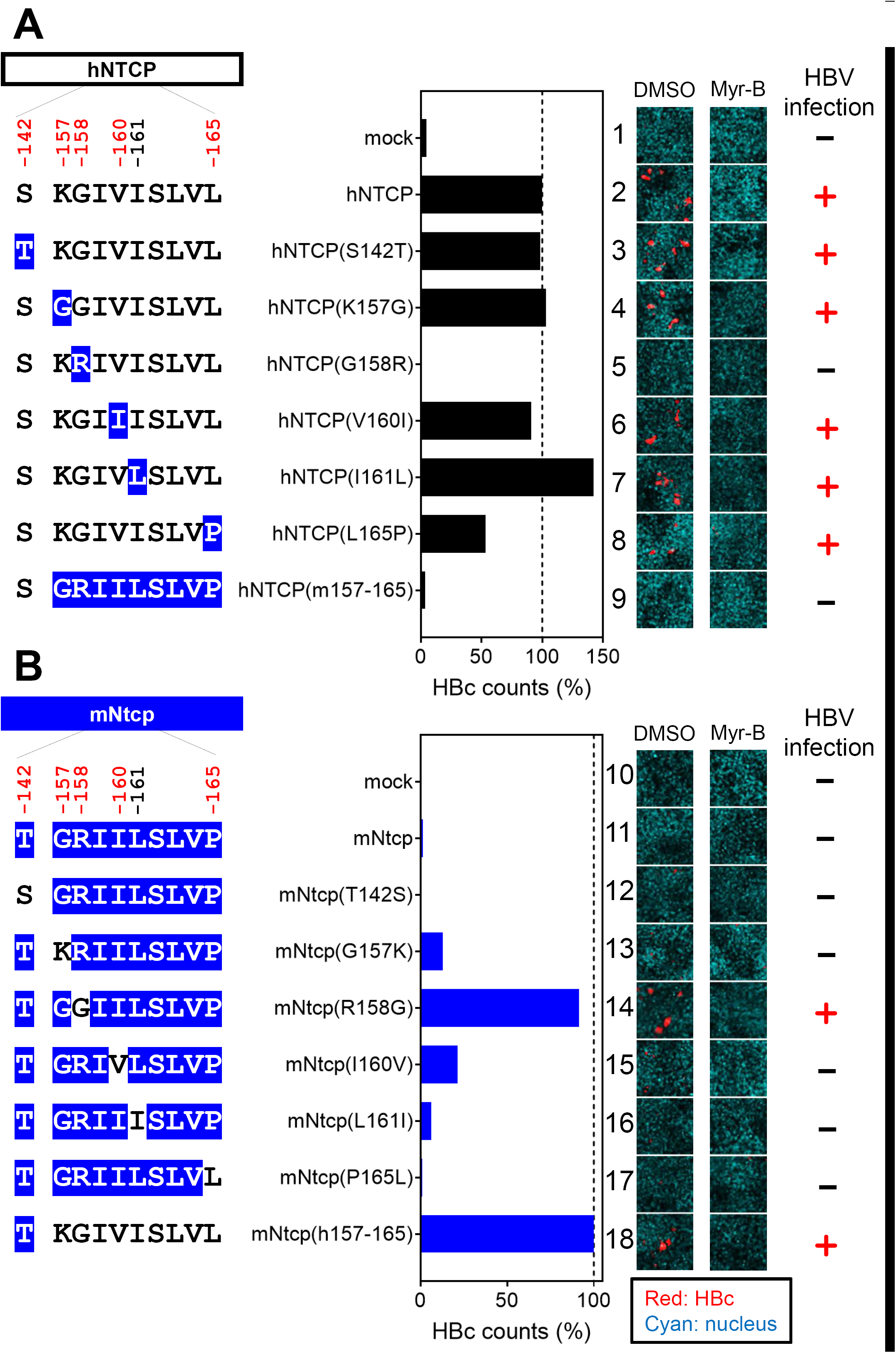
A single positive selection site on NTCP, at aa 158, is a key determinant for supporting HBV infection. Left, Schematic representation of the wild-type and variants of hNTCP (A) and mNtcp (B) (black: human-type residues; blue: macaque-type residues). Middle graphs, HBV susceptibility in HepG2 cells expressing each NTCP construct was determined by detecting HBc-positive cells; the detected value was normalized to the NTCP expression level in the same cell line. Right panels, HBc (red) and the nucleus (cyan) were detected in the HBV infection assay by immunofluorescence analysis in the presence or absence of Myr-B, an HBV entry inhibitor. On the right, the inability or ability to support HBV infection (without Myr-B) for each construct is summarized with-or +, respectively.

We then performed a complementary experiment using the mNtcp backbone and substituting single amino acids with the human residues for the corresponding sites; the resulting mutant mNtcps were examined to determine whether any of the aa substitutions endowed cells with viral entry. Using the same methods as those used in Figure 3A and B, we confirmed the protein expression as well as the transporter activity of the wild-type and mutant mNtcps. Notably, all of the mNtcp derivatives were expressed in HepG2 cells and provided functional bile acid uptake (to levels at least 50 % that of the wild type, with activity 7-to 13-fold higher than the background level) (Figure 3C and D). The HBV infection assay indicated that the wild-type mNtcp did not support HBV infection (Figure 4B, lane 11), while the substitution of the human aa 157-165 in mNtcp [mNtcp(h157-165)] permitted infection (Figure 4B, lane 18), as previously reported (26). Among the single-aa changes, only the substitution to the human sequence at aa 158 (R158G) was sufficient to endow mNtcp with HBV receptor function equivalent to that of mNtcp(h157-165) (Figure 4B, lane 14). Thus, a single positive selection site in *NTCP*, corresponding to aa158, was a key determinant for HBV susceptibility.

### The positive selection site of NTCP is critical for HBV attachment

NTCP is involved in the specific attachment of HBV to the host cell surface through binding to the preS1 region of the large surface protein (LHB) of HBV (41). To explore the impact of the NTCP substitutions on the interaction with HBV preS1, we examined the attachment of a fluorescence-labeled preS1 peptide (preS1-TAMRA) to target cells expressing the wild-type NTCP or its variants (Figure 5A and B, right pictures, red, “DMSO”). To confirm the specificity of the observed preS1-TAMRA signals, the preS1 binding assay was (as above) performed in parallel in the presence of Myr-B, an attachment inhibitor (40) (Figure 5A and B, right pictures, “Myr-B”). Expression in HepG2 cells of variants based on hNTCP or mNtcp backbones was confirmed by immunofluorescence analysis (Figure 5A and B, right pictures, green). As shown in Figure 5A, while expression of hNTCP supported preS1 attachment, replacement of aa 158 in hNTCP with the macaque-type residue [hNTCP(G158R)] completely abrogated the ability to bind to the preS1 peptide (Figure 5A, lane 5). On the other hand, replacement of aa 158 in mNtcp with the human residue [mNtcp(R158G)] endowed mNtcp with preS1 binding activity (Figure 5B, lane 14). Notably, although the preS1 binding activity of mNtcp(R158G) was still approximately 40% that of mNtcp(h157-165), this activity was significantly higher than that of wild-type mNtcp (Figure 5B, lane 11), and this result correlated well with the profile of the HBV infection (Figure 4).

**Figure 5.**
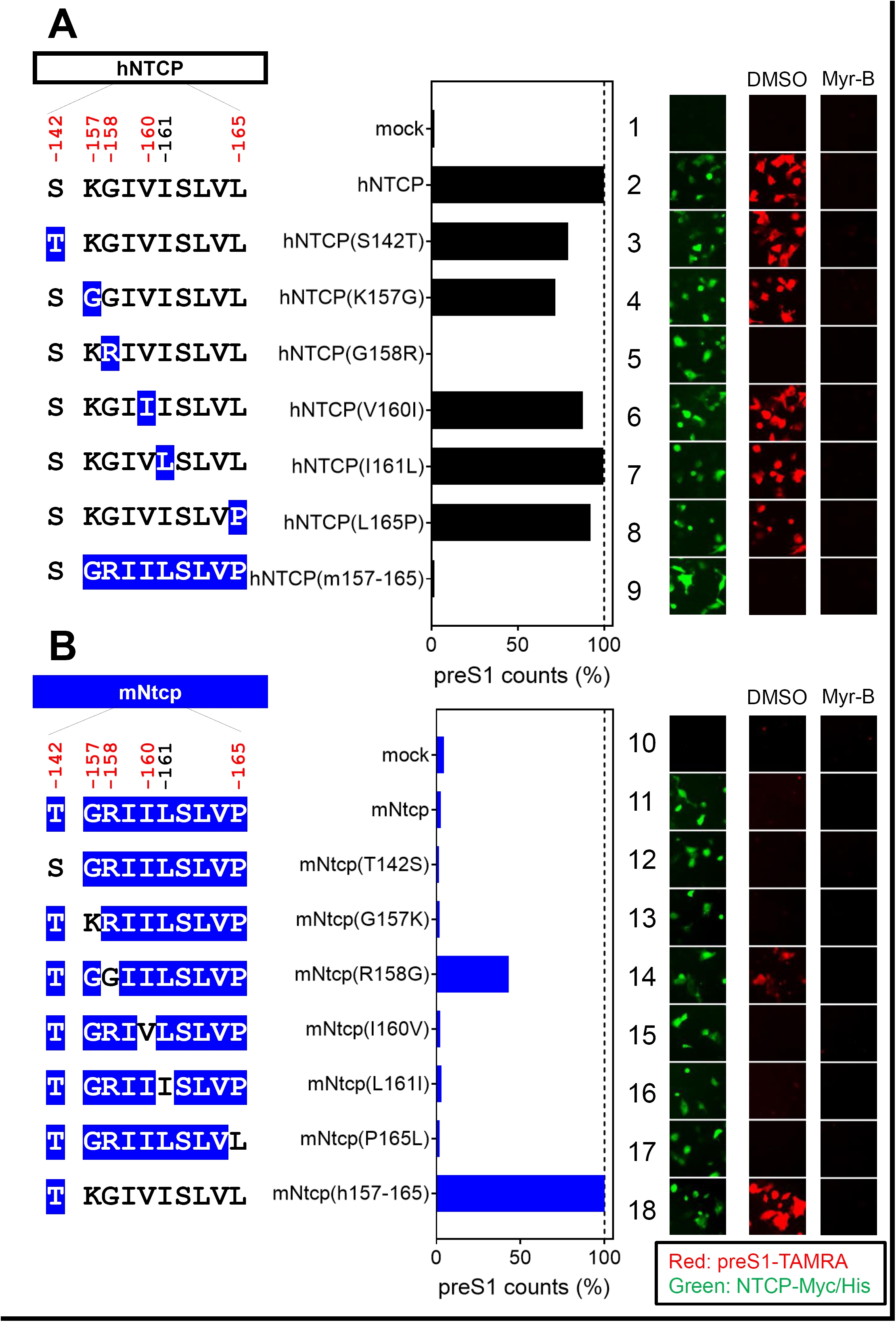
A single positive selection site on NTCP is critical for HBV envelope binding. Left, Schematic representation of hNTCP (A) or mNtcp (B) variants (black: human-type residues, blue: macaque-type residues). Middle graphs, preS1-cell binding mediated by NTCP mutants. The relative amounts of preS1-cell binding were determined by counting preS1-attached cells (red, right panels), and the values were normalized to NTCP expression levels (green, right panels) in the respective cell lines. Right panels, preS1 peptide-cell binding was examined by preS1 binding assay (red) either in the presence or absence of Myr-B. Expression levels of Myc/His-tagged NTCP were simultaneously visualized by immunofluorescence analysis (green).

Hepatitis D virus (HDV) virions harbor an envelope identical or very similar to that of HBV particles, entering cells in a manner similar to that employed by HBV (42). Consistent with the above results obtained in the HBV infection and preS1-binding assays, HDV susceptibility in hNTCP-expressing cells was abrogated by the G158R substitution in hNTCP (Figure 6A, lane 5), whereas the R158G substitution in mNtcp endowed cells with susceptibility to HDV infection (Figure 6B, lane 14). Taken together, these observations suggested that a single position on NTCP, at aa 158, determined the susceptibility to HBV/HDV infection by facilitating viral attachment.

**Figure 6.**
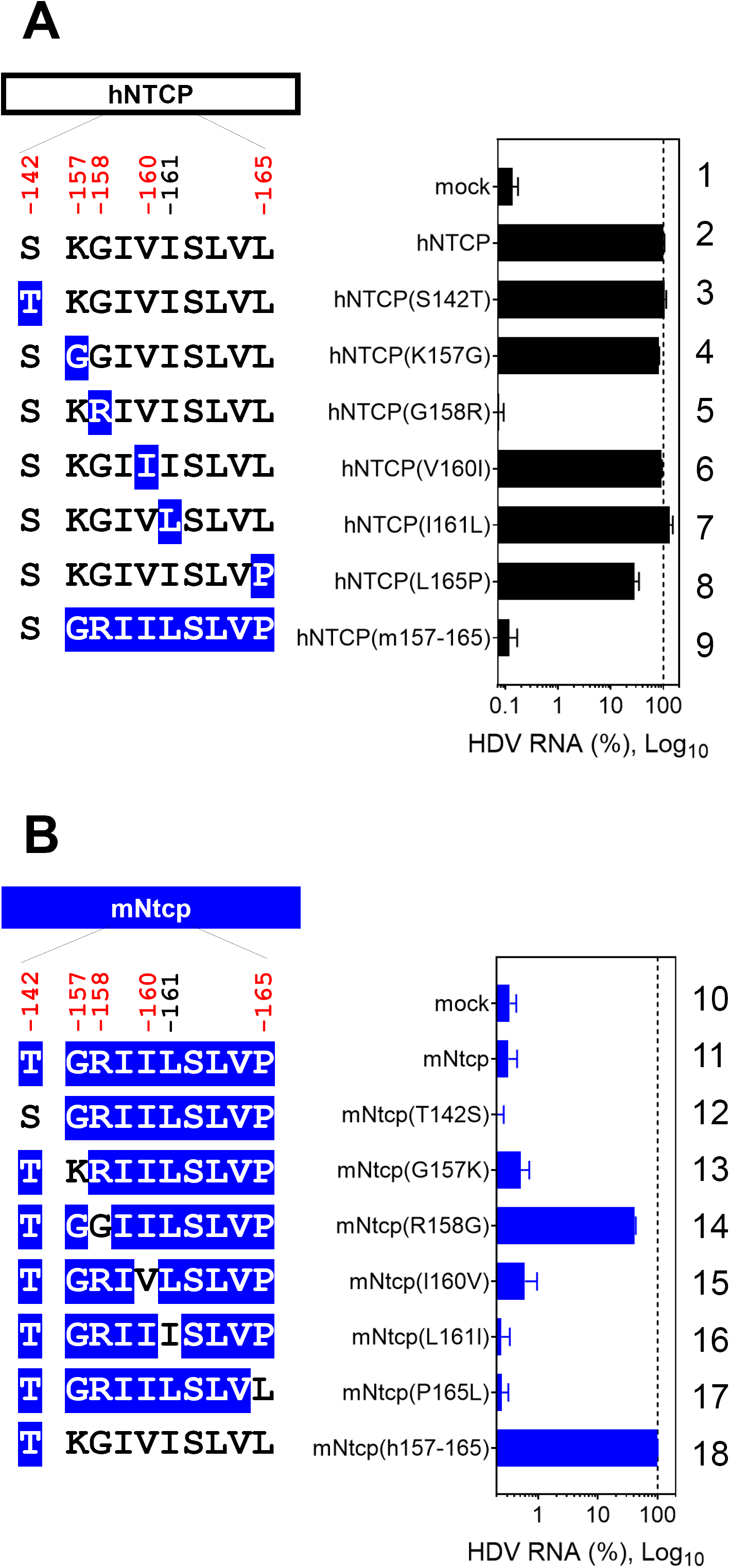
A single positive selection site in NTCP determines HDV susceptibility. Left, schematic representation of hNTCP (A) or mNtcp (B) variants (black: human-type residues; blue: macaque-type residues). Right graphs, HDV infection assay for determining the susceptibility to HDV infection of HepG2 cells expressing each NTCP. Intracellular HDV RNA was normalized by the NTCP expression level in the respective cell line. Results are presented as mean ± SEM (n=3).

## Discussion

Our evolutionary analyses revealed that *NTCP* is highly conserved in mammals (across 66.2 % of the codons), indicating that NTCP is critical for (or advantageous to) host viability. This is in agreement with a previous report showing that *NTCP*-knockout mice could exhibit body weight loss and elevate serum bile acid levels (43). While the majority of *NTCP* sequence was subjected to negative selection, several positive selection sites were inferred. This observation is consistent with analyses of two other virus receptors, TfR1 and NPC1, which showed overall negative selection with a subset of positive selection sites corresponding to the virus-binding domain (31, 32). To verify the biological significance of the positive selection sites in NTCP, we focused on a comparison of the protein between human and the Old-World monkey lineage. Notably, the latter family is phylogenetically the closest to human but Old-World monkeys are not susceptible to HBV infection (38), rendering these simians as the best counterpart for comparison. Interestingly, our infection experiments suggested that a single positively selected site inferred in the present study, aa 158, is a key determinant of HBV’s host species-specificity. The consistent role of aa 158 has been just reported by a very recent paper (44). This observation is supported by the fact that NTCPs that carry the human-type sequence at aa 158 (e.g., NTCP of chimpanzee, orangutan, New-World monkey, treeshrew, and mouse, Figure 1) is known to interact with HBV preS1 (28, 44-46) (although these species except for chimpanzee and treeshrew are not susceptible to HBV infection because of yet unknown restriction mechanism in post-entry steps (44, 46)). Given that the crystal structure of NTCP has not been solved, the exact mechanism whereby aa 158 facilitates HBV entry remains unknown. However, our data suggest that this residue is a primary mediator of the interaction between the preS1 region of the HBV envelope protein and the cell surface (Figure 5). This inference would be analogous to the analysis of TfR1 and NPC1, of which positively selected sites have been suggested to be the interface of virus binding, as a signature of coevolutionary relationship with virus infection (31, 32).

To date, most of the relevant studies have reported that error-prone RNA viruses and lethal viruses (e.g., HIV and filovirus) exert selective pressure on their host proteins (15-22, 32). On the other hand, if a virus is not pathogenic to the host, adaptive mutations in host proteins would not be expected to occur (11). In most cases, HBV infection to adult human is controlled by immunity; only in a minority of cases does chronic infection occur for developing chronic liver disease, and a limited cases develop fulminant hepatitis (47). Therefore, it has been unclear whether HBV could serve as a driver of adaptive evolution in host proteins. However, hepadnavirus infection in malnourished wildlife species, which can be immunocompromised, may results in higher morbidity and mortality (48). In addition, an analysis by Enard *et al.* of the evolutionary patterns of ∼9,900 proteins conserved in mammalian genomes revealed that host proteins that have been in contact with viruses for extended evolutionary periods in a wide range of lineages display higher rates of adaptation to viruses; among these proteins, those that interact with HBV proteins had the strongest signature of adaptation, suggesting the coevolution of host factors and hepadnaviruses (12). However, Enard *et al.* did not provide a molecular analysis showing the detailed gene sites that were under positive selection by virus infection. Therefore, the present study provides the first clear evidence for positive selection (in NTCP) mediated by hepadnaviruses. We hypothesize that the predecessors of Old-World monkeys may have been infected with pathogenic hepadnaviruses that imposed strong selective pressures; while these ancient Old-World monkeys declined or became extinct, their descendants (the extant Old-World monkeys) presumably evolved to escape hepadnavirus infection by accruing a mutation at NTCP aa 158. Indeed, examples of endangerment or extinction of species caused by virulent virus infection have been reported (49-51). Further investigation is needed to determine whether any hepadnavirus exists in extant Old-World monkeys, because the number of primate species surveyed for hepadnaviruses infection has been still limited (38). However, all known Old-World monkey NTCPs carry 158R (i.e., HBV-unsusceptible NTCP) (44), consistent with the apparent absence of hepadnavirus in Old-world monkeys. Simian immunodeficiency virus (SIV) is nonpathogenic in its “natural” hosts, while its virus relatives are highly pathogenic to rhesus macaque, who is an experimental host, and human, who only recently have become hosts (52, 53). These observations support the idea that a virus recently acquired from different species possesses high pathogenic properties and may impose selective genetic pressure on the new host; subsequent host adaptation over the course of a long history of infection would render such a virus less pathogenic to the new host (54). Interestingly, hepadnaviruses have an extremely long history of infection of their respective hosts. The present study demonstrates that the host species-specificity of a virus can be determined by a single positively selected site in a host receptor, as evidenced by a comparison of HBV susceptibility between human and Old-World monkeys and correlation with the sequence of the *NTCP* gene. Although there is a possibility that viruses or factors other than hepadnavirus also exerted selective pressure on NTCP, we provide the first evidence that at least hepadnaviruses can be a driver for inducing adaptive mutations in NTCP.

Surprisingly, our study implies that the HBV cross species barrier is secured by a single amino acid change in the virus receptor. Several studies have reported that a single mutation on a virus envelope protein determines species-specificity (55, 56), but our insight is unique in showing that a single mutation in a host receptor plays a key role in determining the host specificity. It is not known whether all hepadnaviruses (or which hepadnaviruses) use NTCP as their entry receptor. However, tent-making bat hepadnavirus (3) and New-World monkey hepadnaviruses (44) are reported to interact with NTCP, suggesting that NTCP functions as a receptor for a wide range of hepadnaviruses (at least, from bat to human). The present analysis suggests that HBV-related hepadnaviruses drive an adaptive evolution in their hosts’ proteins. Our novel insights are expected to improve our understanding of how hepadnaviruses co-evolved with their hosts over evolutionary intervals, thereby gaining strong host species-specificity.

## Materials& Methods

### Molecular evolutionary analyses

Twenty mammalian full-length NTCP protein sequences were collected from GenBank as listed in Table 1 and aligned by using MUSCLE implemented in MEGA7 (57). The resulting alignment was verified manually at the amino acid level. Then, a Maximum Likelihood (ML) tree of the 20 NTCP sequences was reconstructed using PhyML 3.0 (58) with 1,000 bootstrap resamplings, and used for further analysis (Figure 1). The best fitting substitution model was determined using Smart Model Selection (SMS) (59) implemented in PhyML 3.0. The Akaike Information Criterion (AIC) selected TN93 + G as the best-fit model. To test positive selection or the presence of sites with ω (dN/dS) > 1 in the evolution of NTCP, we performed two pairs of site-models with CODEML implemented in the PAML v 4.8 (34, 35): the first pair involves employed M1a (neutral, 2 site classes, ω > 1 not allowed) versus M2a (selection, 3 site classes, ω > 1 allowed), and the second pair consisted of M7 (neutral, 10 site classes, ω > 1 not allowed) versus M8 (selection, 11 site classes, ω > 1 allowed). FEL, REL (36), and MEME (37) also were employed to detect positive selection sites through the DATAMONKEY webserver (60). HKY85 was determined as the best-fitting substitution model based on the model selection tool in DATAMONKEY. The following default significant cut-off values were used: p-value < 0.1 for FEL, Bayes factor > 50 for REL, and p-value < 0.1 for MEME.

### Plasmids and cells

Plasmids were constructed to permit expression of human NTCP (hNTCP) or macaque Ntcp (*Macaca fascicularis*, mNtcp) tagged with Myc/His at the C-terminus. Specifically, the open reading frame encoding either hNTCP (NM_003049.3) or mNtcp (NM_001283323.1) was inserted between the KpnI and XbaI sites of pEF4-Myc/His A vector (Thermo Fisher Scientific, Whaltham, MA). Individual mutant constructs were constructed using oligonucleotide-directed mutagenesis (61).

HepG2 cells were cultured as described previously (33). To generate NTCP-expressing cells, HepG2 cells were transfected with each NTCP expression plasmid using Lipofectamine 3000 (Thermo Fisher Scientific) according to the manufacturer’s protocol. At 24 h post-transfection, the cells were trypsinized and transferred into 96-well plates; the replated cells then were subjected to preS1 binding assay or virus infection assay (performed as described below) at 48 h post-transfection.

### Western blotting

Cells were treated first with lysing buffer (1 % NP-40, 150 mM NaCl, 50 mM Tris-HCl pH7.5) and then with 250 U Peptide-N-Glycosidase F (PNGase F) to digest N-linked oligosaccharides from glycoproteins. NTCP expression was examined by immunoblotting using monoclonal anti-c-Myc antibody (Santa Cruz Biotechnology, Dallas, TX) at a 1:3,000 dilution, as described previously (55). Beta-actin was detected as a loading control by using monoclonal anti-β-actin antibody (SIGMA, St. Louis, MO) at a 1:10,000 dilution. The chemiluminescence signals were scanned with C-DiGit (LI-COR Biosciences, Lincoln, NE) and analyzed with Image Studio Lite, ver. 5.2 (LI-COR Biosciences).

### NTCP transporter assay

NTCP transporter activity was measured by incubating cells with [^3^H]-taurocholic acid at 37°C for 15 min in a sodium-containing buffer to allow substrate uptake into the cells. After washing to remove free [^3^H]-taurocholic acid, cells were lysed and intracellular radioactivity was measured using a LSC-6100 liquid scintillation counter (Hitachi-Aloka Medical, Tokyo, Japan) as described previously (61). To measure the background signal, the same assay was performed in a sodium-free buffer, in which the NTCP transporter does not function. To evaluate the transporter activity of equivalent amounts of NTCP protein (normalized NTCP transporter activity), we divided the value for bile acid uptake by the level of NTCP expression determined by immunofluorescence analysis. Results are presented as mean ± SEM (n=3).

### HBV preparation and infection

HBV inocula were derived from the culture supernatant of Hep38.7-Tet cells (genotype D); these inocula were prepared as described previously (62). For the HBV infection assay, HBV was infected into recipient cells at 12,000 genome equivalents (GEq)/cell in the presence of 4 % polyethyleneglycol (PEG) 8000 for 16 h, followed by washing (to remove free virus) and culturing the cells for an additional 12 days, according to the previously described protocol (63). Indirect immunofluorescence analysis for detection of HBc was conducted to evaluate HBV infection; this technique was performed essentially as described previously (64). Briefly, after fixation with 4% paraformaldehyde and permeabilization with 0.3% Triton X-100, the cells were treated with a polyclonal rabbit anti-HBV core antibody (Thermo Fisher Scientific) at a 1:100 dilution, and then with an Alexa Fluor 594 (Thermo Fisher Scientific)-conjugated donkey anti-rabbit IgG secondary antibody at a 1:500 dilution. Stained cells were photographed using a BZ-X710 microscope (Keyence, Osaka, Japan), and the images were analyzed with BZ-X Analyzer 1.3.1.1 (Keyence). The relative number of HBV-infected cells was determined by counting HBc-positive cells using the Hybrid Cell Count module of the Keyence analyzer, and the values were normalized for the NTCP expression levels determined by immunofluorescence analyses.

### preS1 binding assay

To examine binding between the preS1 region in the large surface protein of HBV (LHBs) and host cells, cells were exposed to 20 nM 6-carboxytetramethylrhodamine-labeled preS1 peptide (TAMRA-preS1) at 37 °C for 30 min; unbound peptide then was removed by washing. To simultaneously detect Myc/His-tagged NTCP, the cells were fixed with 4 % paraformaldehyde and permeabilized with 0.3% Triton X-100, and the tagged NTCP was detected with a monoclonal anti-c-Myc antibody at a 1:1,200 dilution and with an Alexa Fluor 488-conjugated goat anti-mouse IgG secondary antibody (Thermo Fisher Scientific) at a 1:1,000 dilution; nuclei were stained with 4’,6-diamidino-2-phenylindole (DAPI) at a 1:5,000 dilution. Fluorescence was quantified as described above.

### HDV preparation and infection

HDV was prepared from culture supernatants of Huh-7 cells transfected with pSVLD3 (kindly provided by Dr. John Taylor at the Fox Chase Cancer Center) and pT7HB2.7 as described previously (65, 66). The cells were inoculated with HDV at 15 GEq/cell in 5 % PEG 8000 for 16 h, followed by washing (to remove free virus) and culturing of the cells for an additional 6 days. Total intracellular RNA was extracted and reverse transcribed using a TaqMan Gene Expression Cells-to-Ct kit (Thermo Fisher Scientific) according to the manufacturer’s protocol. Intracellular HDV RNA was quantified by real-time reverse transcription PCR (RT-PCR) as described previously (67). Results are presented as mean ± SEM (n=3).

## Acknowledgments

HDV expression plasmids were kindly provided by Dr. John Taylor at the Fox Chase Cancer Center. This study was supported by the Japan Society for the Promotion of Science (JSPS) Research Fellow Grant (JP16J40232); KAKENHI (JP17H04085, JP66KT0111, JP17K15708); the JST CREST program; the Japan Agency for Medical Research and Development, AMED (JP18fk0310114j0002, JP18fk0310101j1002, JP18fk0310103j0002, JP18fk0310103j0202, JP18fm0208019j0002, JP18fk0210036j0001, and JP18fk0210009j0003); the Takeda Science Foundation; and the Pharmacological Research Foundation, Tokyo.

